## Supplementary Figures 1-8 for "Inhibitory Gating of Coincidence-Dependent Sensory Binding in Secondary Auditory Cortex"

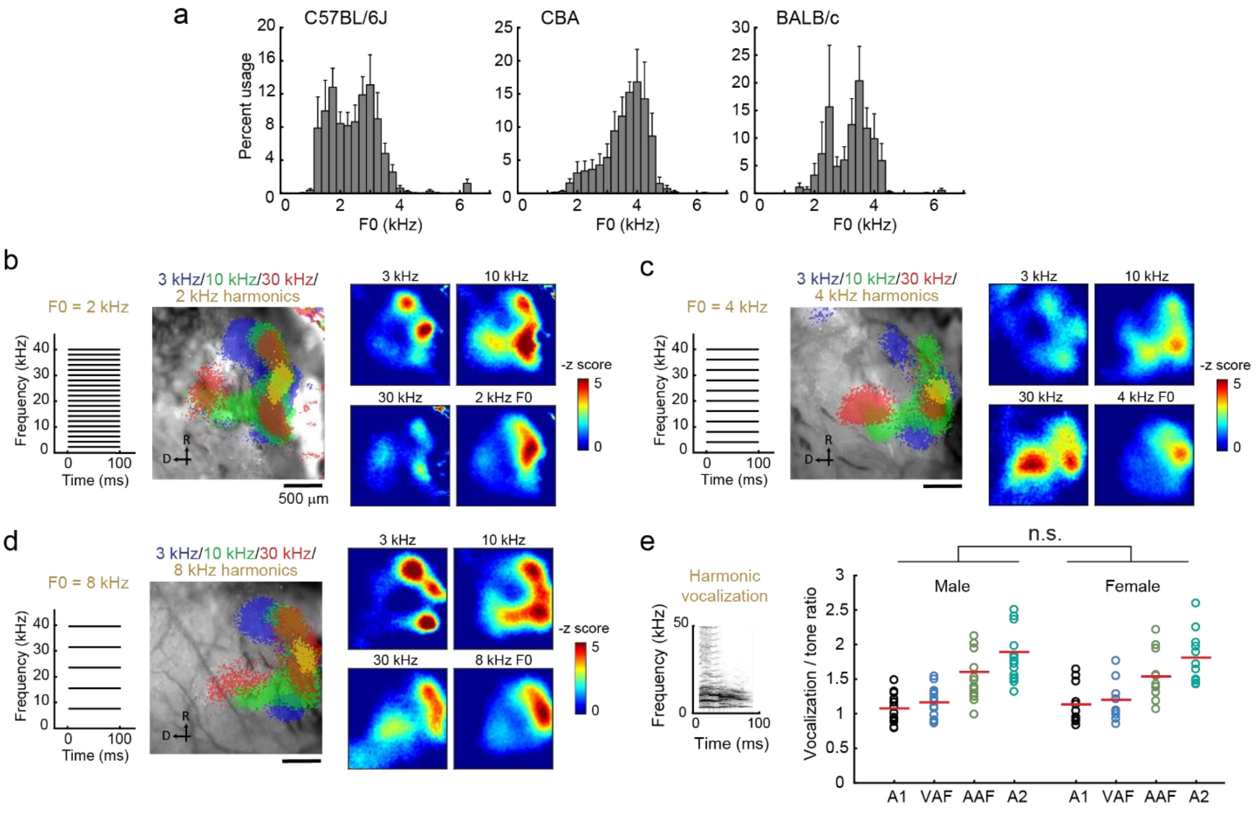


**Extended Data Figure 1. Additional analyses for mapping harmonic responses.**

**(a)** Histograms showing the usage probability of fundamental frequency (F0) for harmonic vocalizations in three strains—left, B6 (n = 5 mice, median: 2.5 kHz, 80% of F0s fell between 1.5 and 3.3 kHz); middle, CBA (n = 3 mice, median: 3.5 kHz, 80% fell between 2.1 and 4.3 kHz); right, BALB/c (n = 4 mice, median: 3.3 kHz, 80% fell between 2.6 and 3.9 kHz). Results are mean ± SEM. **(b)** Left, spectrogram of artificial 2 kHz-F0 harmonics. Middle, thresholded intrinsic imaging signal responses to pure tones as well as harmonics in a representative mouse. Right, heat maps showing z-scored response amplitudes. **(c)** Data for F0 = 4 kHz harmonics. **(d)** Data for F0 = 8 kHz harmonics. **(e)** Ratio of harmonic vocalization to pure tone response amplitudes in each of the auditory cortical areas for male and female mice. Left, males (n = 15 mice); right, females (n = 11 mice). No difference was observed between males and females (male vs female, $p = 0.862$; areas, $p < 0.0001$; interaction, $p = 0.873$; Two-way ANOVA). Red lines show mean.

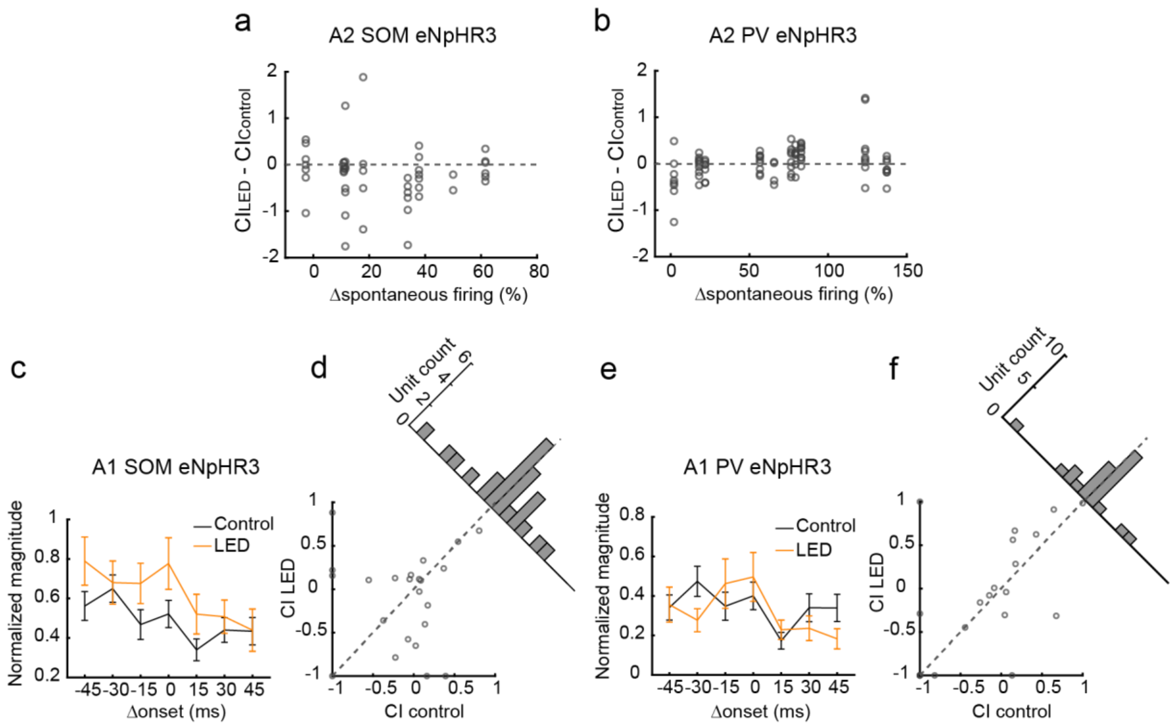


**Extended Data Figure 2. Optogenetic inactivation of inhibitory neurons does not change coincidence preference in A1.**

**(a)** Scatter plot showing change in CI against spontaneous firing rate change triggered by SOM cell photoinactivation in individual regular-spiking units (n = 6 mice, 47 units, allowing duplication of the same units across two photostimulation intensities). Units with spontaneous firing rate less than 0.25 Hz were excluded. **(b)** The same plot for PV cell photoinactivation (n = 7 mice, 81 units). **(c)** Summary plot showing the response amplitudes of A1 regular-spiking units to 4-kHz harmonics with various Δonsets during control and SOM cell inactivation trials. Responses are normalized to the maximum response amplitude in the control condition in each unit and averaged across all units (n = 5 mice, 24 units in the A1 superficial layer). Data are mean ± SEM. **(d)** Scatter plot showing CI during control and SOM cell inactivation trials. The oblique histogram illustrates the changes in CI with LED. p = 0.913 (two-sided paired t-test). **(e)** Summary plot showing the response amplitudes of A1 regular-spiking units during control and PV cell inactivation trials (n = 6 mice, 25 units in the A1 superficial layer). **(f)** Scatter plot showing CI during control and PV cell inactivation trials. The oblique histogram illustrates the changes in CI with LED. p = 0.551.

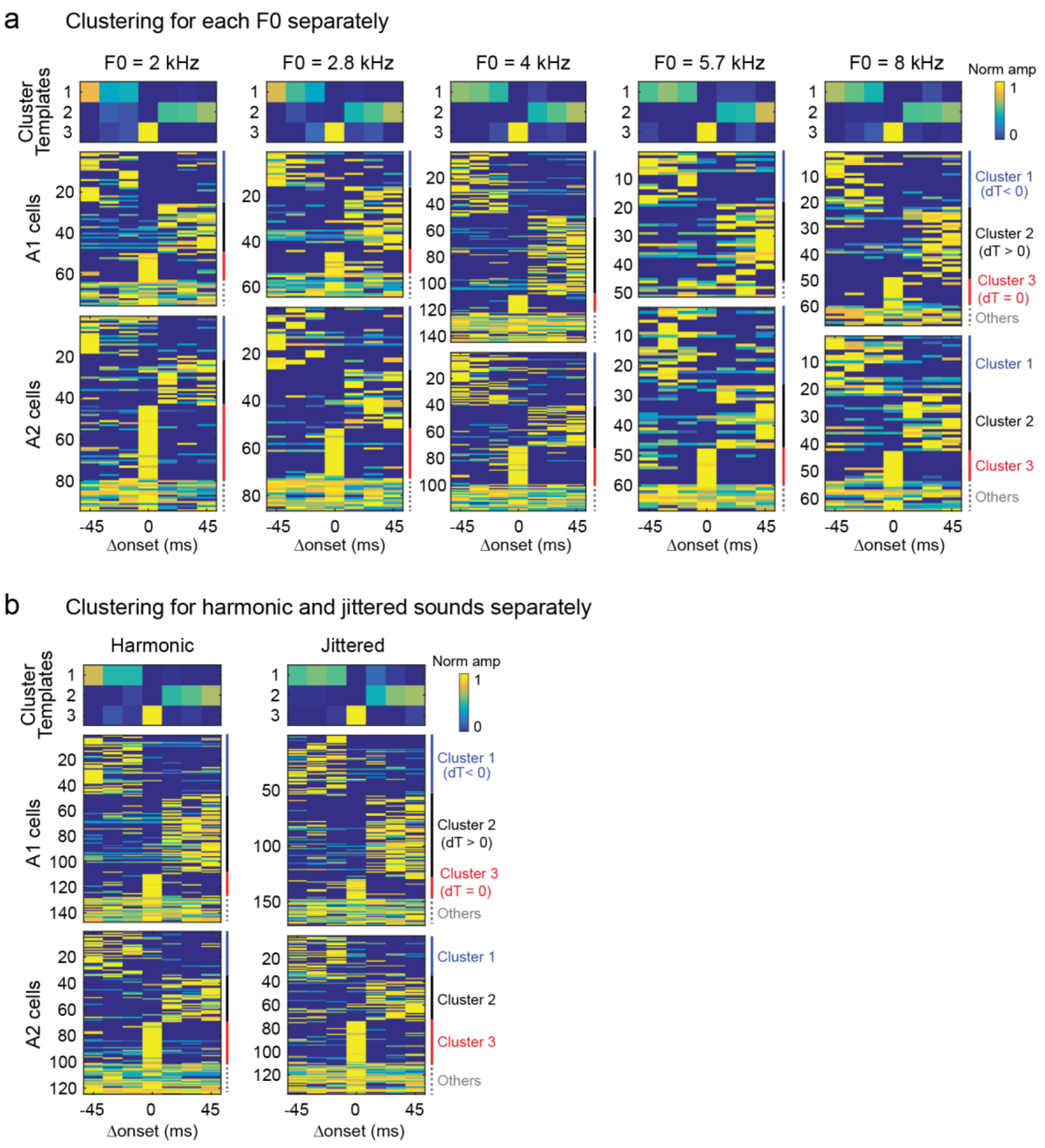


**Extended Data Figure 3. Reproducible clustering of A1 and A2 neurons into negative shift-, positive shift-, and coincidence-preferring groups for individual F0s and jitters.**

**(a)** Clustering data independently conducted for each F0. The order of appearance of three clusters was matched to that in Figures 5 and 6. F0 = 2 kHz (A1: n = 75 cells, A2: n = 94 cells); F0 = 2.8 kHz (A1: n = 64 cells, A2: n = 86 cells); F0 = 4 kHz (A1: n = 147 cells, A2: n = 119 cells); F0 = 5.7 kHz (A1: n = 51 cells, A2: n = 68 cells); F0 = 8 kHz (A1: n = 67 cells, A2: n = 64 cells). Clustering was robust even when conducted separately for individual F0s. **(b)**

Clustering data independently conducted for harmonic and jittered sounds, Harmonic (A1: n = 147 cells, A2: n = 124 cells); Jittered (A1: n = 171 cells, A2: n = 136 cells).

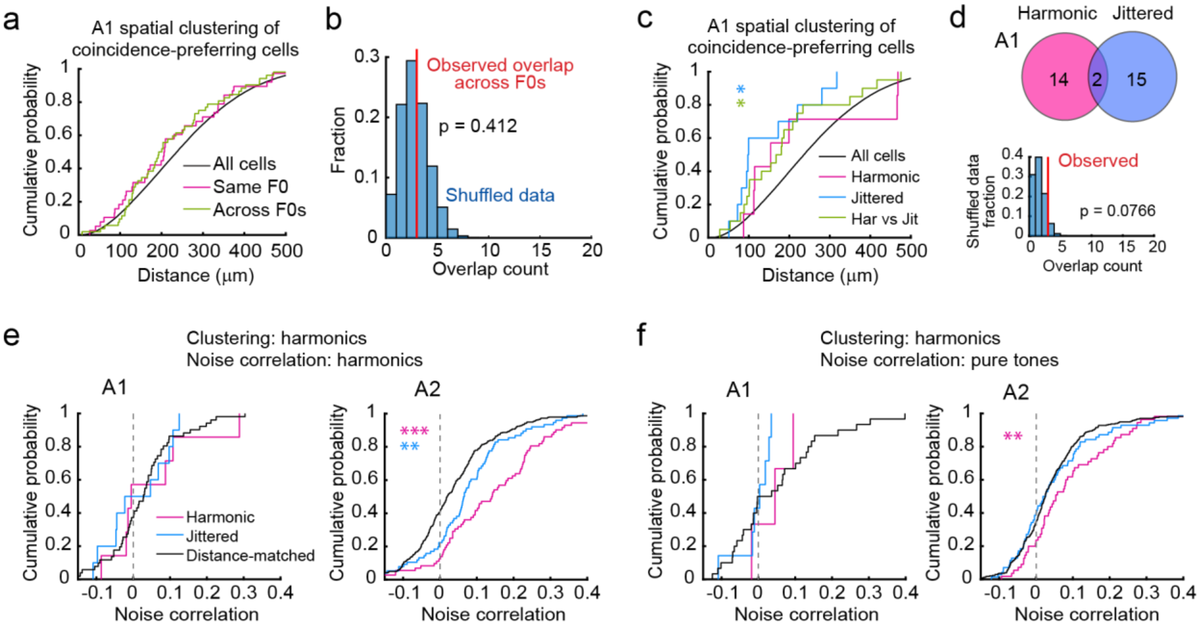


**Extended Data Figure 4. Lack of spatial clustering or functional subnetwork for coincident harmonics-preferring neurons in A1.**

**(a)** Cumulative probability plot of spatial distance between all cells (black), between coincidence-preferring cells with the same F0 (magenta), and between coincidence-preferring cells across F0s (green) in A1 (n = 46662, 38, 52 for All cells, Same F0, and Across F0s). **(b)** Fraction of overlap between coincidence-preferring cells across F0s for A1. Red line, observed overlap count; histogram, distribution of overlap count for shuffled data (10,000 repetitions). **(c)** Cumulative probability plot of spatial distance in A1 between all cells (black), between coincidence-preferring cells for harmonics (magenta), between coincidence-preferring cells for jittered sounds (blue), and between coincidence-preferring cells for harmonic and jittered sounds (green) (n = 44610, 7, 10, 20 for All cells, Harmonic, Jittered, and Har vs Jit). *$p < 0.05$ (Wilcoxon rank sum test). **(d)** Top, Venn diagram showing the overlap between coincidence-preferring clusters for harmonic and jittered sounds in A1. Bottom, observed overlap of coincidence-preferring cells for harmonics and jittered sounds (red line) compared to shuffled data (histogram, 10,000 repetitions). **(e)** Cumulative probability plots of noise correlation between coincidence-preferring cells for harmonic and jittered sounds in A1 and A2. Data for A2 is the same as Figure 6i (A1: n = 7, 10, 51; A2: n = 72, 76, 399 for Harmonic, Jittered, and Distance-matched). **$p < 0.01$, ***$p < 0.0001$ **(f)** Cumulative probability plots of noise correlation between coincidence-preferring cell pairs during presentation of pure tones (A1: n = 3, 7, 30; A2: n = 55, 70, 330 for Harmonic, Jittered, and Distance-matched).

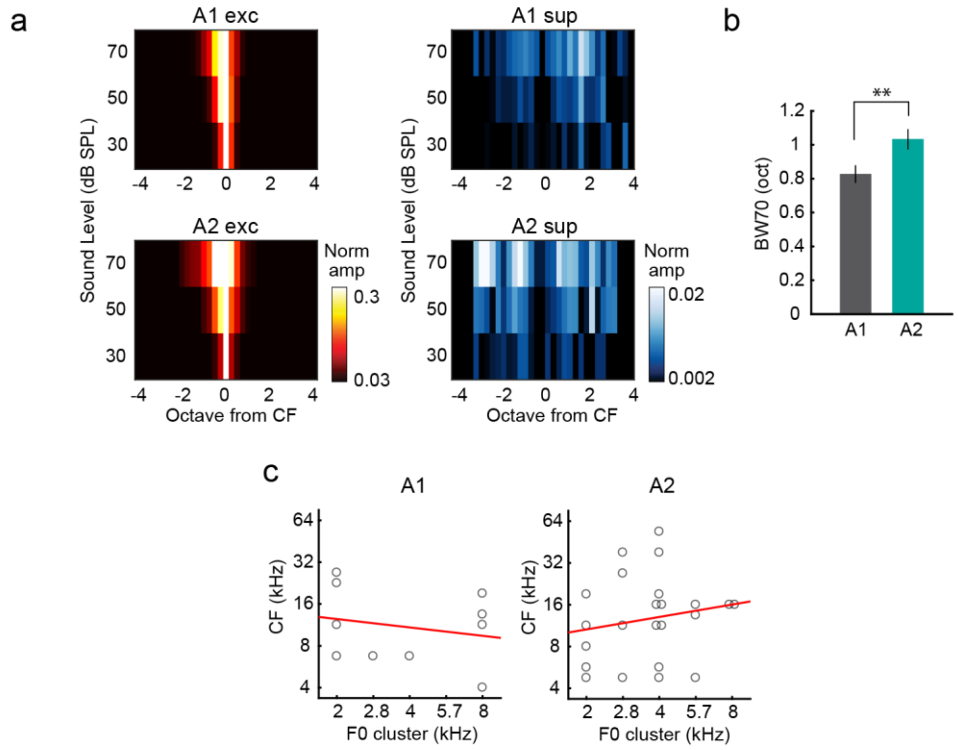


**Extended Data Figure 5. Tonal receptive fields of A1 and A2 neurons.**

**(a)** Tonal receptive fields of excitatory and suppressive responses to pure tones averaged across all cells with excitatory responses in A1 (n = 296 cells, 11 mice) and A2 (n = 232 cells, 10 mice). Responses are centered around the characteristic frequency of excitatory response for each cell. **(b)** Tuning broadness of neurons in each area measured as bandwidth at 70 dB SPL (BW70). **p < 0.01 (two-sided t-test). Results show mean ± SEM. **(c)** Characteristic frequency of coincidence-preferring neurons for each F0 in A1 and A2 (A1: slope = -0.216, p = 0.549, n = 10 cells responsive to pure tones out of 26 cells which were classified as coincidence-preferring; A2: slope = 0.178, p = 0.418, n = 23 out of 69 cells). We did not observe a relationship between preferred F0 and characteristic frequency of coincident harmonics-preferring neurons, suggesting that these neurons do not encode pitch. Red lines show linear regression.

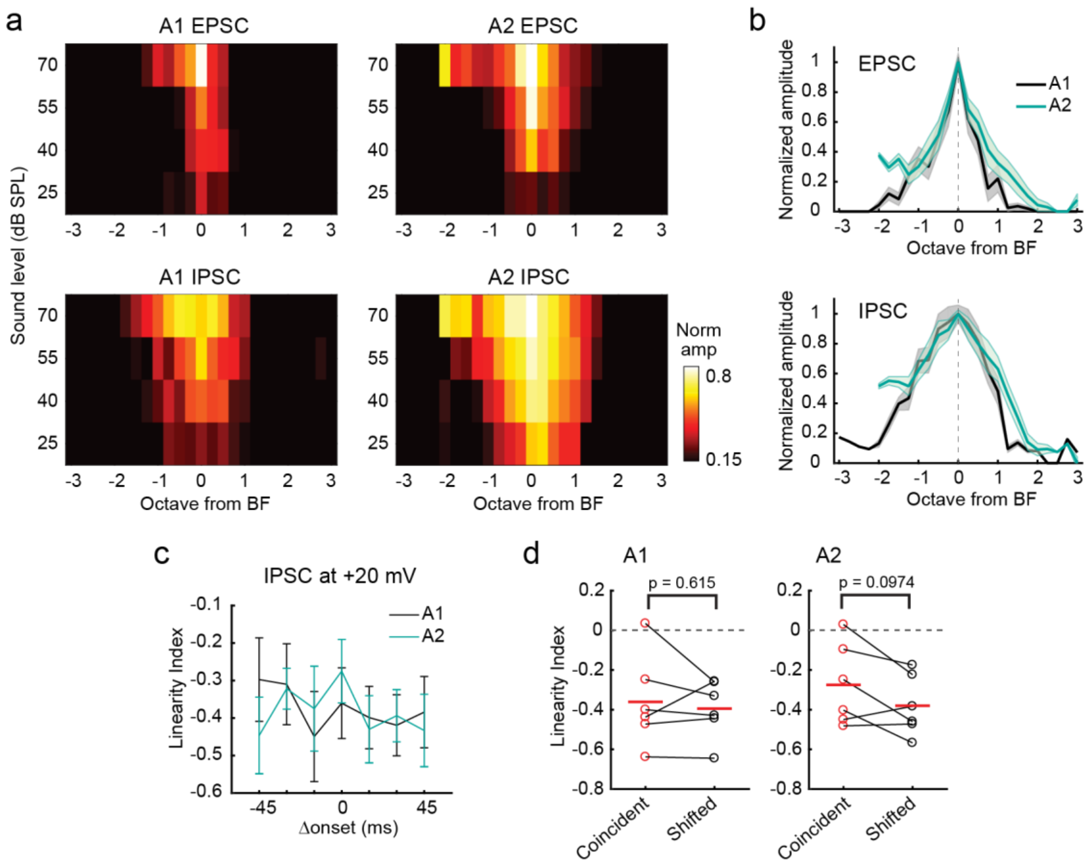


**Extended Data Figure 6. Tonal receptive fields and three-tone harmonics responses of synaptic currents in A1 and A2.**

**(a)** Tonal receptive fields of EPSCs and IPSCs in response to pure tones averaged across cells in A1 (EPSCs: n = 16 cells, IPSCs: n = 16 cells) and A2 (EPSCs: n = 6 cells, IPSCs: n = 6 cells). Responses are centered around the best frequency of excitation for each cell. **(b)** Summary of EPSC and IPSC frequency tuning in A1 and A2. Response amplitudes are normalized to their individual peaks. Dark line, mean; shading, SEM. **(c)** Linearity index of IPSCs calculated for each Δonset in A1 and A2 (A1: n = 6; A2: n = 6 cells). Results show mean ± SEM. **(d)** Summary plots showing the linearity index of IPSCs calculated for coincident and shifted harmonics (A1: p = 0.615; A2: p = 0.0974, paired t-test). Red lines show mean.

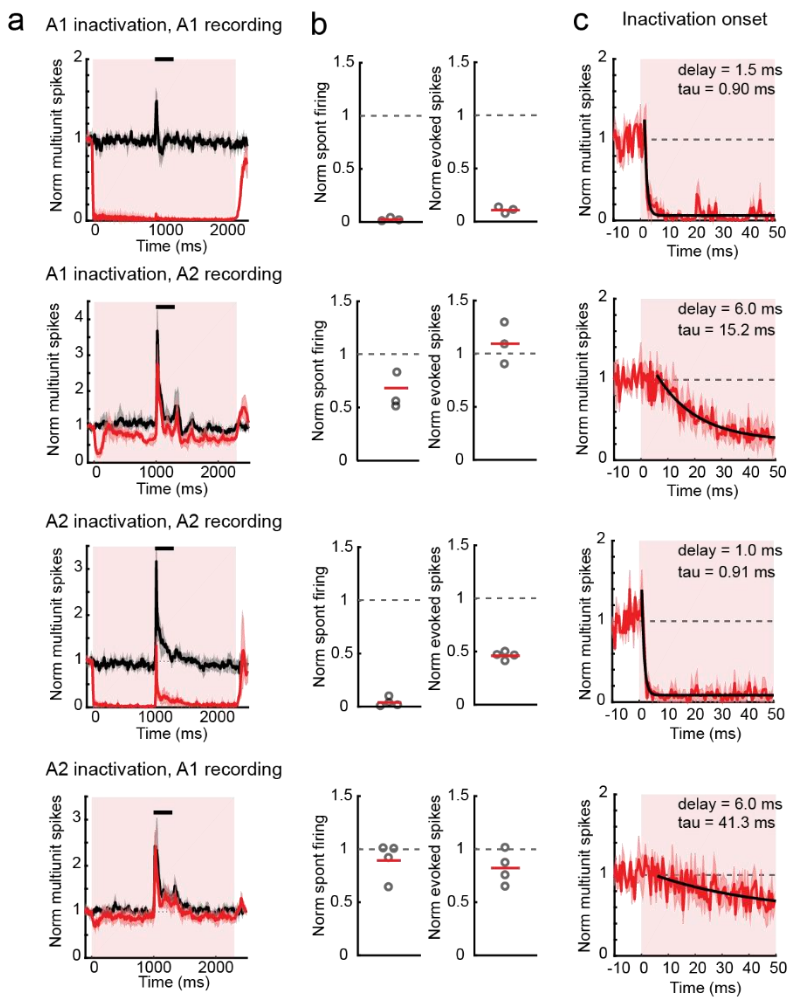


**Extended Data Figure 7. Optogenetic inactivation is restricted to the targeted areas.** **(a)** Functional inactivation of cortical areas across recording conditions: A1 inactivation, A1 recording (n = 3 mice); A1 inactivation, A2 recording (n = 3); A2 inactivation, A2 recording (n = 4); A2 inactivation, A1 recording (n = 4). PSTHs show multiunit spikes during control (black) and photostimulation (red) trials. Solid lines and shades show mean ± SEM. **(b)** Summary results showing the reduction of spontaneous and evoked firing rate during photostimulation. Red lines show mean. **(c)** Multiunit spikes during the first 50 ms of photostimulation showing inactivation kinetics rapidly after LED onset. Black lines, single-exponential fit. A1 inactivation, A1 recording and A2 inactivation, A2 recording data display rapid decay time constants with < 1 ms latency, indicating their direct inactivation. In contrast, A1 inactivation, A2 recording and A2 inactivation, A1 recording data show only slow decay with long latency, confirming the lack of direct photoinactivation.

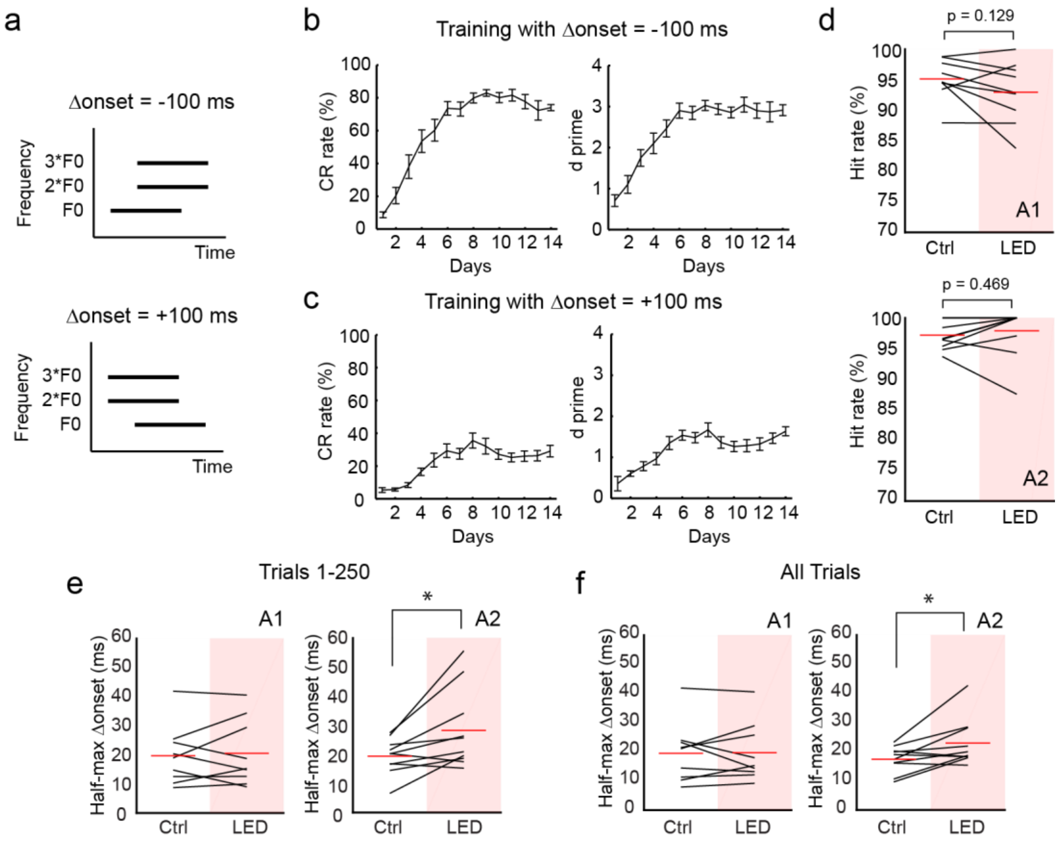


**Extended Data Figure 8. Harmonics discrimination training with negative and positive onset shifts.**

**(a)** Schematic for three-tone harmonic stimulus with -100 ms and +100 ms Δonsets. **(b)** Learning curves averaged across all tested mice (n = 19) during training with Δonset of -100 ms. Left, average correct rejection rate over days. Right, average d prime over days. Results are mean ±SEM. (The same data as Fig. 8c) **(c)** Same as (b) except for Δonset of +100 ms. **(d)** Top, hit rate for target coincident harmonics with and without A1 inactivation (n = 9 mice). Bottom, hit rate for target coincident harmonics with and without A2 inactivation (n = 10 mice). **(e)** Half-max Δonsets with and without inactivation of A1 (left) or A2 (right), using the first 250 trials (A1: n = 9; A2: n = 10 mice; the same data as Fig. 8h). **(g)** The same data using all trials in each mouse (486 +/- 31 trials). A1 control: 19.5 ± 3.4, LED: 19.7 ± 3.4; A2 control: 17.4 ± 1.4, LED: 23.0 ± 2.6. *p = 0.0273 (two-sided Wilcoxon signed rank test).
